# GPRC6A is a Potential Therapeutic Target for Metformin Regulation of Glucose Homeostasis in Mice

**DOI:** 10.1101/2024.08.19.608635

**Authors:** Min Pi, Rupesh Agarwal, Micholas Dean Smith, Jeremy C. Smith, L. Darryl Quarles

## Abstract

Understanding the mechanism of metformin actions in treating type 2 diabetes is limited by an incomplete knowledge of the specific protein targets mediating its metabolic effects. Metformin has structural similarities to L-Arginine (2-amino-5-guanidinopentanoic acid), which is a ligand for GPRC6A, a Family C G-protein coupled receptor that regulates energy metabolism. Ligand activation of GPRC6A results in lowering of blood glucose and other metabolic changes resembling the therapeutic effect of metformin. In the current study, we tested if metformin activates GPRC6A. We used Alphafold2 to develop a structural model for L-Arginine (L-Arg) binding to the extracellu-lar bilobed venus flytrap domain (VFT) of GPRC6A. We found that metformin docked to the site in the VFT that overlaps the binding site for L-Arg. Metformin resulted in a dose-dependent stimulation of GPRC6A activity in HEK-293 cells transfected with full-length wild-type GPRC6A but not in untransfected control cells. In addition, metformin failed to activate an alternatively spliced GPRC6A isoform lacking the putative binding site in the VFT. More specifically, mutation of the predicted metformin key binding residues Glu170 and Asp303 in the GPRC6A VFT resulted in loss of metformin receptor activation in vitro. The in vivo role of GPRC6A in mediating the effects of metformin was tested in Gprc6a-/- mice. Administration of therapeutic doses of metformin lowered blood glucose levels following a glucose tolerance test in wild-type but not Gprc6a-/- mice. Finally, we EN300, created by adding a carboxymethyl group from L-Arg to the biguanide backbone of metformin. EN300 showed dose-dependent stimulation of GPRC6A activity in vitro with greater potency than L-Arginine, but less than met-formin. Thus, we suggest that GPRC6A is a potential molecular target for metformin which may be used to understand the therapeutic actions of metformin and develop novel small molecules to treat T2D.

## Introduction

Metformin (1,1-dimethylbiguanide) is a common treatment for type 2 diabetes (T2D)(1, 2). Metformin has many metabolic effects (3), including lowering of blood glucose through associated inhibition of hepatic gluconeogenesis, enhanced peripheral insulin responsiveness, and stimulation of glucose uptake in skeletal muscle, as well as facilitating weight loss and lower of free fatty acid (FFA levels) in association with suppression of adipose lipolysis. Metformin treatment is also associated with beneficial cardiovascular, anti-aging(4), and anticancer effects(5).

The molecular mechanisms whereby metformin regulates glucose and energy metabolism are poorly understood. At the cellular level, metformin inhibits complex I in the mitochondrial respiratory chain (6). Metformin activates AMPK by upstream liver kinase B1 (LBK1), increases AMP/ATP ratio by inhibition of mitochondrial respiratory chain complexes and reduces lactate and glycerol metabolism to glucose by inhibiting mitochondrial glycerophosphate dehydrogenase(7). In addition, via LKB1-AMP-mTOR pathway, metformin increases autophagy leading to improved mitochondrial function. This drug also has AMPK-independent effects (1). Metformin inhibits glycerol-3-phosphate dehydrogenase and alters of cellular redox balance via inhibition of the glycerophosphate shuttle leading to impaired hepatic gluconeogenesis. These cellular effects are purported to be direct and presupposes the cellular uptake of metformin through OCT1 mediated organic cation transport (8–10). Metformin also activates mitogen-activated protein kinase (MAPK) and other signaling pathways, including SIRT1, FOXO, NFκβ, and DICER1(4, 11). In addition, metformin stimulates the release of several hormones that regulate glucose metabolism, including FGF21, Glp-1, testosterone, and adiponectin (10, 12–14).

Metformin’s pleiotropic effects are postulated to be mediated by its interaction with a wide array of protein targets and pathway networks (15). Molecular docking analysis has identified metformin binding to AMPK(16), Cyclin E1 (CCNE1) a proto-oncogene (17), and the epigenic modifiers H3K27me3 demethylase KDM6A/UTX (18). It is unusual for a drug with specific therapeutic effects to target so many structurally diverse proteins, and moreover, none of these targets fully explain metformin’s effects on glucose and energy metabolism. Also, metformin is thought to function intracellularly(8), but whether metformin might activate a cell surface GPCR has not been previously explored.

GPRC6A is a member of the Family C G-protein coupled receptors that regulates energy metabolism (19–22). GPRC6A is expressed in key metabolic organs, including pancreatic β-cells, skeletal muscle and liver, where genetically engineered loss-of-function mouse models show that GPRC6A regulates glucose, amino acid and fat metabolism (20, 23–50). Using Gprc6aflox/flox mice to selectively delete *Gprc6a* in pancreatic β-cells, skeletal muscle, hepatocytes and adipocytes, loss-of GPRC6A has been respectively shown to decrease insulin secretion and β-cell proliferation (20, 51), lower muscle glucose uptake (52, 53), stimulate hepatic glucose and fatty acid metabolism (54, 55) and suppresses lipolytic activity in adipocytes(56). In addition, ligand activation of GPRC6A stimulates insulin secretion and beta-cell proliferation in the pancreas (21), fibroblast growth factor 21 (FGF-21) release and enhances glucose and fat metabolism in liver hepatocytes (54), stimulates interleukin 6 (IL-6) secretion and glucose utilization in skeletal muscle myocytes (57). GPRC6A activation also induces adiponectin release and lipolytic activity in adipocytes in white fat (56), stimulates testosterone production from testicular Leydig (54, 58), and augments glucagon-like peptide 1 (GLP-1) secretion from gastrointestinal enterocytes (59, 60). Many of these effects overlap those of metformin on energy metabolism.

GPRC6A is structurally characterized by an extracellular “venous fly trap (VFT)” region connected to a heptahelical transmembrane (7-TM) domain. GPRC6A is activated by several orthosteric ligands, including basic L-amino acids (such as L-Lys, L-Arg, and L-ornithine) and divalent cations (such as Ca^2+^ and Mg^2+^). Also, positive allosteric modulators (PAMs) of GPRC6A include osteocalcin, testosterone, NPS-568, and di-phenyl and tri-phenyl compounds (61, 62).

Insights into structural basis for GPRC6A interaction with multiple ligands is revealed by high resolution structural data of the related mGluR5 (63) and CaSR (64–66). These studies show that mGluR5 and CaSR have similar orthosteric binding site for L-glutamate and L-tyrosine in the VFT cleft, and two positive modulating allosteric (PAM) sites, one located in the VFT for peptides and the other in the 7TM for small molecules (63–66). Similarly, an Alphafold2 model of GPRC6A predicts that L-amino acids (such as L-Lys, L-Arg, and L-ornithine) and divalent cations (such as Ca^2+^ and Mg^2+^) bind to a putative orthosteric binding site in the VFT extracellular domain, and that the peptide osteocalcin, a positive allosteric modulators of GPRC6A binds to a site in the VFT (61). This structural model also predicts another PAM site in the 7TM for testosterone and di-phenyl and triphenyl compounds (61, 62).

Interestingly, there are chemical similarities between the guanidine group in metformin (C_4_H_11_N_5_) and the molecular features of the GPRC6A ligand L-Arginine (C_5_H_14_N_4_O_2_) that may allow molecular interactions with the same protein target.

Because of the overlapping physiological effects of metformin and GPRC6A activation on glucose and energy metabolism and chemical similarities between metformin and L-Arg, we tested if metformin binds to and activates GPRC6A and examined metformin’s glucose lowering effects in Gprc6a^-/-^ mice.

## Materials and Methods

### Structure and binding site modeling

The previous work laid the foundation of this work, where we developed (AlphaFold2-derived) 3D models of GPCR6A has been shown to be useful in predicting the interactions of osteocalcin with GPCR6A(67). The top predicted structure was then aligned with two established Class C GPCRs, the calcium-sensing receptor (CaSR) bound to L-Tryptophan (PDB ID: 7DTU) (68)and the metabotropic glutamate receptor 5 (mGluR5) bound to the orthosteric ligand L-quisqualate (PDB ID: 6N51)(63). This alignment revealed a conserved orthsteric pocket in GPRC6A isoform 1, positioned similarly to the L-Trp binding site in CaSR. Based on this structural homology, we modeled L-Arginine in the putative binding site of GPRC6A. We took the aligned structures of mGluR5 bound to L-Trp bound to position an L-Arg residues into the corresponding site in the GPCR6A model to generate a bound pose. The side chain of the pockets were relaxed using induced-fit docking and molecular-mechanics (MM) based ligand-receptor geometry optimization (see below). All alignments and structure editing was performed using the CCG Molecular Operating Environment (MOE) (Chemical Computing Group, Montreal, QC).

### Induced Fit Molecular Docking

Docking of ligands was performed in two phases-phase 1: refinement of L-Arg positioning and phase 2: docking of metformin and EN300 via templated induced fit docking and subsequent MM-geometry optimization. In phase 1, induced fit docking was performed in MOE with the internal ‘induced-fit’ module with the GBVI/WSA dG scoring function. The all poses were then visually inspected for an evaluated for reasonable ligand-protein contact distances and the top pose containing the maximum number of predicted ligand-protein interactions and was followed by an MM-based energy minimization/geometry optimization with a convergence criteria of 0.1 kcal/(mol * Å^2^). Energy minimization was performed using the Amber10-EHT force-field and the R-field solvation approach. Interaction maps were then obtained from the energy minimized system. For the second phase of docking of metformin and EN300, the template-docking protocol from MOE was used with the refined L-Arg position used as the template. Following docking, the top pose was then visualized and followed by MM energy minimization and the energy minimized poses were used to obtain interaction maps. Energy minimization of EN300 and metformin were performed with the same settings as L-Arg. These docked poses were used to suggest mutations for experimental validation.

### Reagents

L-Arginine and EN300 were ordered from MilliporeSigma (11009 and ENAH31552054; St. Louis, MO, USA). Metformin was purchased from Thermo Fisher Scientific (M2009; Waltham, MA, USA).

### Measurement of Total and Phospho-ERK by ERK Elisa Analysis

All culture reagents were from Invitrogen (Waltham, MA, USA). Human embryonic kidney HEK-293 cells were obtained from American Type Culture Collection and cultured in DMEM medium supplemented with 10% fetal bovine serum and 1% Penicillin/Streptomycin (P/S). Briefly, HEK-293 cells transfected with/without human GPRC6A isoforms or mutant cDNA plasmids were starved by overnight incubation in serum-free DMEM/F12 containing 0.1% bovine serum albumin (BSA) and stimulated with various ligands at different doses. ERK activation was assessed 20 min after treatment by using ERK1/2 (phospho-T203/Y204) ELISA Kit (Invitrogen) corrected for the amount of total ERK using ERK1/2 (Total) ELISA Kit (Invitrogen) to measure ERK levels.

### Measurement of cAMP accumulation

HEK-293 transfected with human GPRC6A isoform or mutant cDNA plasmids cells (10^5^ cells/well) (69) were cultured in 24-well plates in DMEM supplemented with 10% fetal bovine serum and 1% penicillin/streptomycin (100 U/mL of penicillin and 100 μg/mL of streptomycin) for 48 hours followed by 4 hours incubation in DMEM/F12 containing 0.1% BSA and 0.5 mM IBMX to achieve quiescence. Quiescent cells were treated with vehicle control, various ligands at concentration as indicated for 40 minutes at 37°C. Then, the reaction was stopped, and the cells lysed with 0.5 mL 0.1N HCl. cAMP levels were measured by using Cyclic AMP EIA kit (Cayman Chemical) following the manufacture’s protocol.

### Site-directed mutagenesis

*In-vitro* mutagenesis by PCR-mediated recombination QuikChange II XL Site-Directed Mutagenesis Kit (Agilent) was performed using the cDNA of human GPRC6A cloned in the vector pcDNA3 (Invitrogen) as template. The primer sets as following. E170A.sen: GGGTTATGCATCAACTGCAGAAATCCTGAGTGA; E170A.antisen: GTTGATGCATAACCCACCTGTGGCATGA; D303A.sen: TGCTAGTGCTAATTGGTCAACTGCCACCA; and D303A.antisen: CAATTAGCACTAGCAATCCACATCTTATTT. All mutants were confirmed by sequencing.

### Mouse models and glucose tolerance test

The *Gprc6a^-/-^* mouse model was created by replacing exon 2 of the GPRC6A gene with the hygromycin resistance gene, as described previously (23). Mice were maintained and used in accordance with recommendations as described (National Research Council 1985; Guide for the Care and Use of Laboratory Animals Department of Health and Human Services Publication NIH 86–23, Institute on Laboratory Animal Resources, Rockville, MD) and following guidelines established by the University of Tennessee Health Science Center Institutional Animal Care and Use Committee. The animal study protocol was approved by the institutional review boards at University of Tennessee Health Science Center Institutional Animal Care and Use Committee.

8-week-old wild-type (*Gprc6a^+/+^*) and global knock out (*Gprc6a^-/-^*) 8-week-old female mice were fed normal chow (2018; Harlan Teklad, Madison, WI, USA) or high fat diet (06414; Harlan Teklad, Madison, WI, USA) for 4 weeks, then mice were injected intraperitoneal with 4 mg/g metformin body weight, or vehicle (saline; 10 μl/g body weight). Glucose tolerance test (GTT) was assessed in 6 h fasted mice after i.p injection of 2 g/kg body weight glucose. Tail vein blood glucose was subsequently measured using a handheld glucometer (Accy-Chek, Roche Diagnostics, San Diego, CA, USA) at baseline and after 15, 30, 60, 90 and 120 min (57, 70). Area under the curve (AUC) for glucose was calculated using GraphPad Prism 5.

### Statistics

We evaluated differences between groups with the Student’s *t* test, and for multiple groups by two-way ANOVA, followed by *a post-hoc* Tukey’s test. Significance was set at pL<L0.05. All values are expressed as means ± SEM. All computations were performed using the Statgraphic statistical graphics system (STSC Inc., Rockville, MD, USA).

## Results

### L-Arginine is the orthosteric ligand for GPRC6A

To understand the structural basis for amino acid sensing in GPRC6A, we created a model based on the closely related calcium sensing receptor CASR and mGlu (see method for details). Cryo-EM data show that CASR binds the amino acid L-Tyrosine at the orthosteric pocket of the large extracellular Venus fly trap (VFT) motif to induce a conformational change from open inactive to an active closed configuration. Calcium binds to other sites in the VFT of CASR, resulting in additional conformational changes to fully activate the receptor, thereby imparting its functions as a calcium sensing GPCR.

L-Arginine, which is composed of guanidine, amino and carbolic acid groups, activates GPRC6A (Figure 1 A). To explore L-Arginine binding to GPRC6A we developed a homology model using Alpahfold2 and assessed if L-Arginine binds to a homologous orthosteric ligand binding site located in the VFT of GPRC6A., analogous to L-Tyrosine binding to CASR. We found that the guanidium group can form multiple hydrogen bonds (Figure 1 A and B). In our model L-Arginine is predicted to bind to the orthosteric pocket in the VFT domain, forming contacts with residues Tyr148, Glu170, Gln280, Asp303 and Asn304 (Figure 1B). L-Arginine molecule interacts with Glu170 and Asp303 via hydrogen bonds. Glu170 is encoded by exon 2, and Asp is encoded by exon 3 (the region that lost in GPRC6A isoform 2, see Figure 4A below).

**Figure 1.**
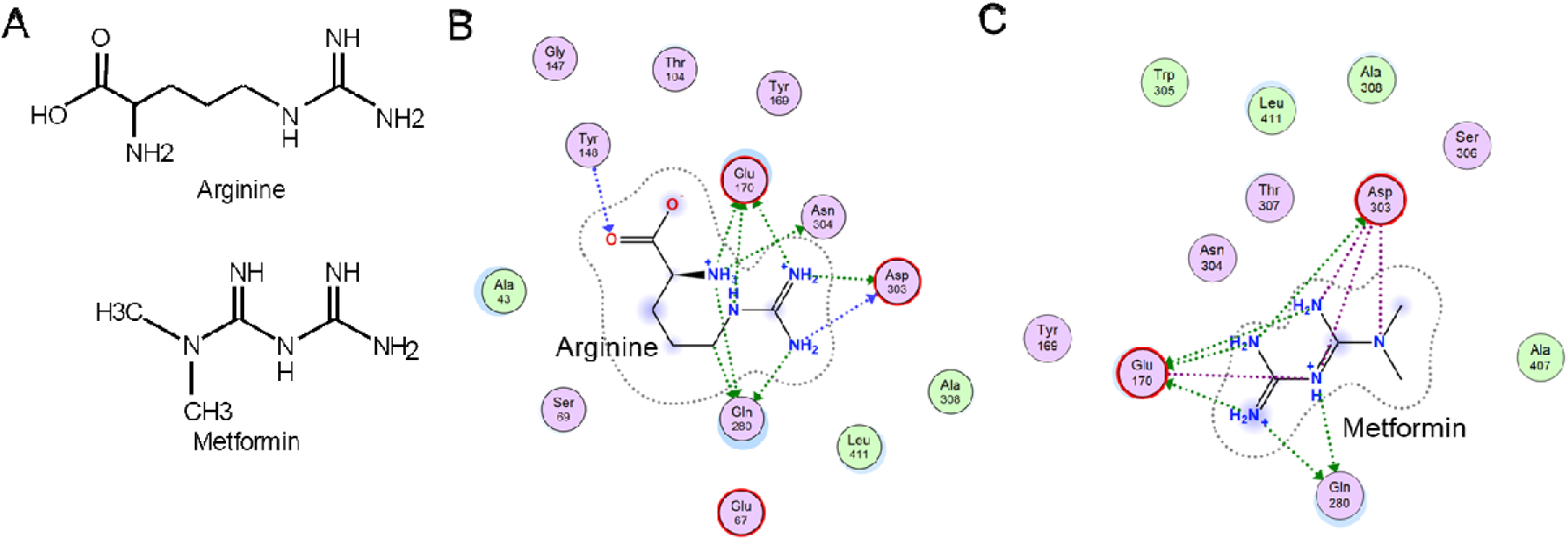
A. Comparison of metformin and L-arginine chemical structures. B and C. 2D ligand binding interaction maps of L-Arginine (B) and metformin (C) docking to the VFT of GPRC6A.

### Metformin is an orthosteric ligand for GPRC6A

Metformin is a biguanide that has chemical similarity to the guanidine group in L-Arginine (Figure 1A). To determine if GPRC6A could be a target for metformin, we docked metformin to our structural model of GPRC6A and found that the “metformin-like” core of L-Arginine has very similar contacts with metformin. In the 2D ligand-protein interaction maps, Glu170 appears to be an important residue for binding, and Gln280 and Asp303 may help hold metformin in place for guanidium groups. Metformin is predicted to bind to residues Ala170 and Asp303 in the region predicted to bind L-Arginine (Figure 1B and C). Additional rendering of metformin binding site shows that metformin docks to the pocket formed between the “top” and “bottom” lobes of the VFT and away from the VFT dimer interface (Figure 2), data derived from performing a protonation state assignment calculation and an energy minimization of the ligand and the pocket using the molecular operating environment (MOE) program. The relaxed docked pose of metformin is consistent with the L-Arginine in the orthosteric pocket.

**Figure 2.**
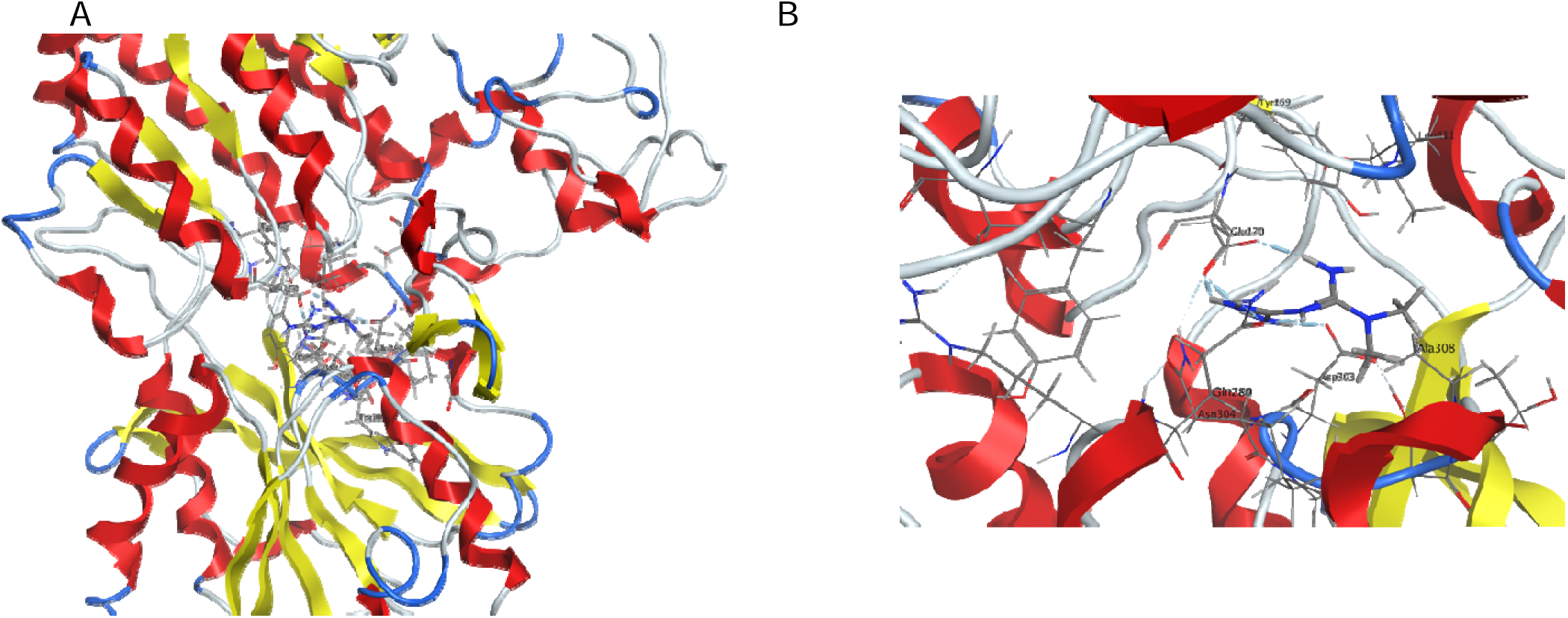
Rendering of metformin binding to the predicted L-Arg binding site in the VFT of GPRC6A. A. A “zoom-out” showing the relative position of the metformin with respect to a rendering of the monomeric VFT. B. A “zoom-in” image of the binding site, with interacting residues labeled. This shows metformin docks to the pocket formed between the “top” and “bottom” lobes of the VFT and away from the VFT dimer interface.

In contrast, docking of L-Tryptophan that is the amino acid ligand for the related receptor CaSR into GPRC6A that the amino group on the backbone orients towards the pocket, but the rest of the L-Tryptophan is exposed, suggesting that it won’t fit in the pocket (Supplemental data: Figure S1C, Table S1).

### Metformin activates GPRC6A signaling *in vitro*

Next, we compared the ability of L-Arginine and metformin to activate full-length and mutant GPRC6A *in vitro*. Consistent with prior reports, we found that L-Arginine dose-dependently stimulate ERK activity and cAMP accumulation in the presence of 1 mM calcium (Figure 3A and B). L-Arginine had no effect in cell not transfected with GPRC6A (Figure 3A and B). The EC50 for L-Arginine stimulated ERK phosphorylation and cAMP accumulation are 5.13 or 15.07 mM. L-Arginine activation of GPRC6A *in vivo* would be in the presence of serum ionized calcium in the1.05-1.3 mmol/L range and circulating levels of L-Arginine are reported range from 72 to 114 μM in humans (71). Our data suggests that super-physiological levels of L-Arginine may be required to activate GPRC6A *in vivo* (50).

**Figure 3.**
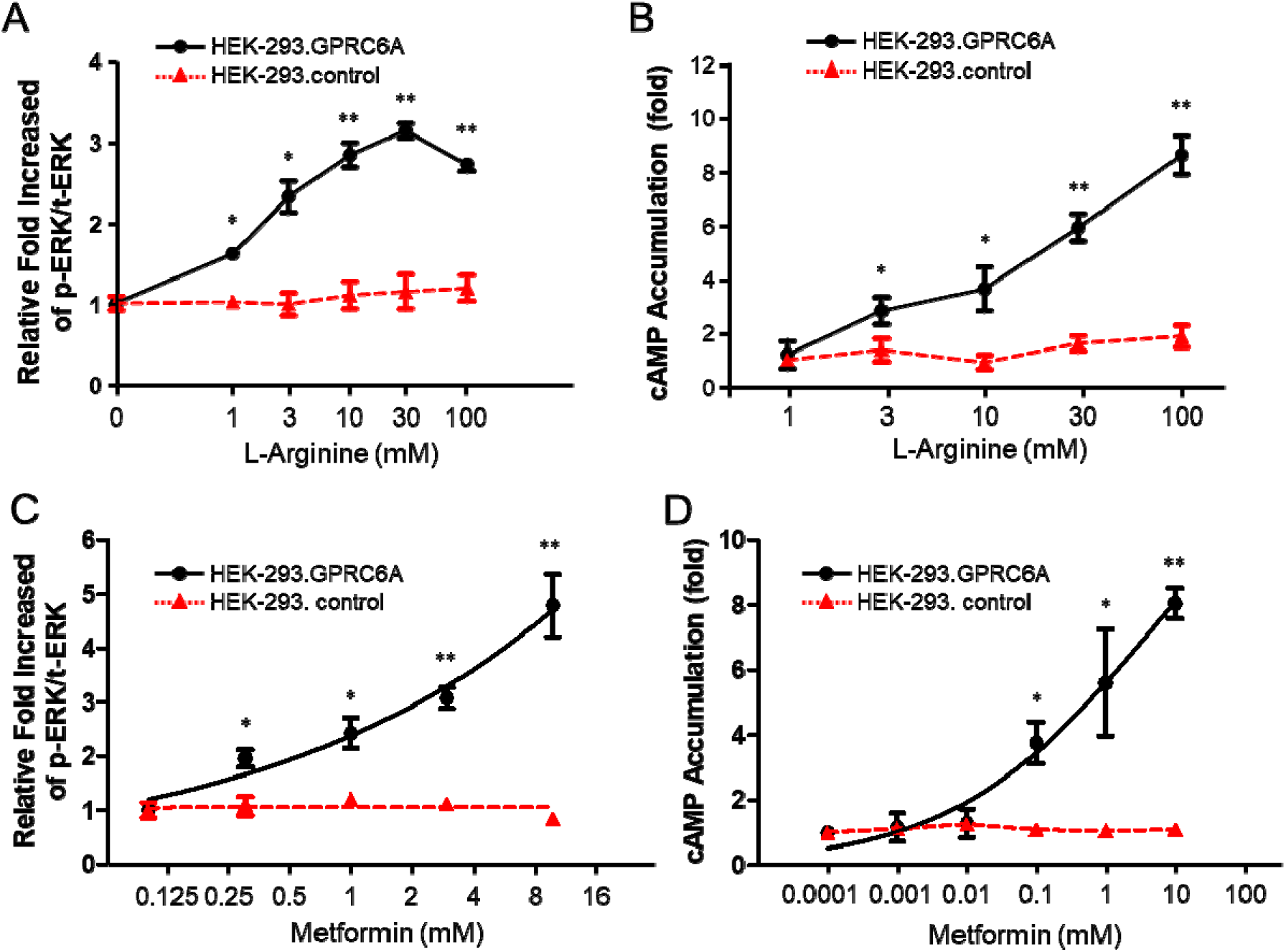
*In vitro* assessment of GPRC6A activation. A and B. L-Arginine dose-dependent stimulates GPRC6A-mediated ERK phosphorylation (A) and cAMP accumulation (B) in HEK-293 cells transfected with GPRC6A but not in non-transfected cells. C and D. Dose-dependent effects of metformin on GPRC6A-mediated ERK activation (C) and cAMP accumulation (D). HEK-293 cells were transfected with cDNA plasmids of GPRC6A for 48 hours, after incubated in Dulbecco’s modified Eagle’s medium/F-12 containing 0.1% bovine serum albumin quiescence media for 4 hours, then exposed to L-Arginine or or metformin at indicated concentrations for 15 minutes for ERK activation, or 40 minutes for cAMP accumulation details as described under “Methods”. Results are means + SE. *P<0.05, **P<0.001 that indicates a significant difference from control and stimulation groups.

Consistent with the predicted binding, we found that metformin resulted in a dose-dependent activation ERK and cAMP signaling in HEK-293 cells transfected with GPRC6A, but not in untransfected controls (Figure 3C and D). The EC50 for metformin stimulated ERK phosphorylation and cAMP accumulation are 0.534 or 0.284 mM. As a point of reference, circulating metformin concentrations after a single therapeutic dose range between 10 and 40 micromolar (72), suggesting GPRC6A could potentially be a clinical target for high doses of metformin.

### Glu170 and Asp303 in VFT of GPRC6A are important residues for L-Arginine and metformin binding

We confirmed the importance of the VFT and orthsteric pocket in metformin binding by using isoforms of GPRC6A which have deletions in the VFT domain due to alternative splicing. Isoform 2, which is missing a portion of Exon 3 encoding Asp303 (D303) the VFT (Figure 4A), lost metformin stimulation of ERK and cAMP (Figure 4B and C). To further assess the prediction, we mutated two key residues in GPRC6A orthosteric pocket, E170A and D303A. The mutation to either of these resulted in the loss of metformin stimulation of ERK and cAMP activity in transfected cells confirming the binding pocket and the key interactions (Figure 4D and E).

**Figure 4.**
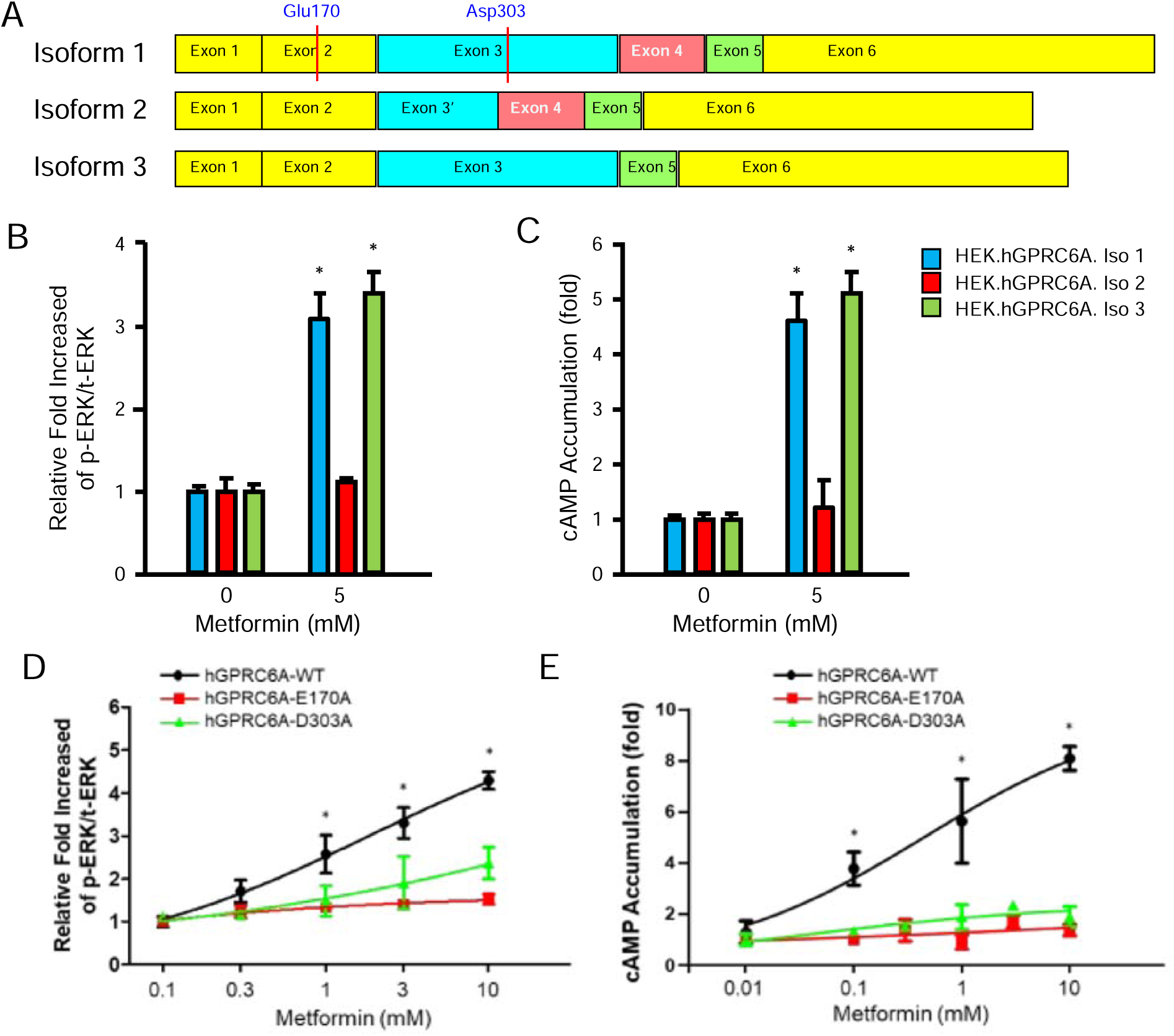
Isoforms and mutants of GPRC6A attenuated activities on response to metformin. a. Placement of exons and its location in GPRC6A isoform 1, 2 and 3 structure. The positions of metformin orthosteric binding pocket residues in GPRC6A are indicated by blue literals. Comparison of dose-dependent effects of metformin on GPRC6A isoforms mediated ERK activation (B) and cAMP accumulation (C). HEK-293 cells were transfected with cDNA plasmids of isoform 1, 2 and 3 GPRC6A for 48 hours, after incubated in Dulbecco’s modified Eagle’s medium/F-12 containing 0.1% bovine serum albumin quiescence media for 4 hours, then exposed to L-Arg or Ocn at indicated concentrations for 15 minutes for ERK activation, or 40 minutes for cAMP accumulation details as described under “Methods”. Comparison of dose-dependent effects of metformin on ERK activation (D) and cAMP accumulation (E) in HEK-293 cells transfected with the cDNA plasmids of WT, E170A, or D303A hGPRC6A. HEK-293 cells were transfected with cDNA plasmids of of WT, E170A, or D303A GPRC6A for 48 hours, after incubated in Dulbecco’s modified Eagle’s medium/F-12 containing 0.1% bovine serum albumin quiescence media for 4 hours, then exposed to metformin at indicated concentrations for 15 minutes for ERK activation, or 40 minutes for cAMP accumulation details as described under “Methods”. *P<0.05 that indicates a significant difference from control and stimulation groups.

Since L-Arginine and metformin are predicted to bind to overlapping sites, we examined the effects of combining L-Arginine and metformin in our GPRC6A activation assays. 10 and 20 mM L-Arginine resulted in a respective 4 and 6 fold increase in ERK activity in HEK-293 cells expressing GPRC6A. The addition of metformin reduced L-Arginine stimulation of GPRC6A, consistent with competitive binding to the same pocket.

### EN300 is a ligand for GPRC6A

To further test the hypothesis that both the carboxyl and guanidine groups are important in GPRC6A activation, we tested a related compound EN300. EN300 has a carboxyl group added to the metformin biguanide structure (Figure 5A). EN300 resulted in a dose-dependent activation of GPRC6A transfected into HEK293 cells (Figure 5B and C). The EC50 for EN300 stimulated ERK phosphorylation and cAMP accumulation are 2.331 or 3.448 mM, respectively. The EC50 for EN300 was greater than L-Arginine, but less than metformin, suggesting that the additional carboxyl group decreased the ligand potency. Docking of EN300 (SI Figure 2) to the same pocket revealed a similar binding pose to metformin and L-Arg (Figure S2).

**Figure 5.**
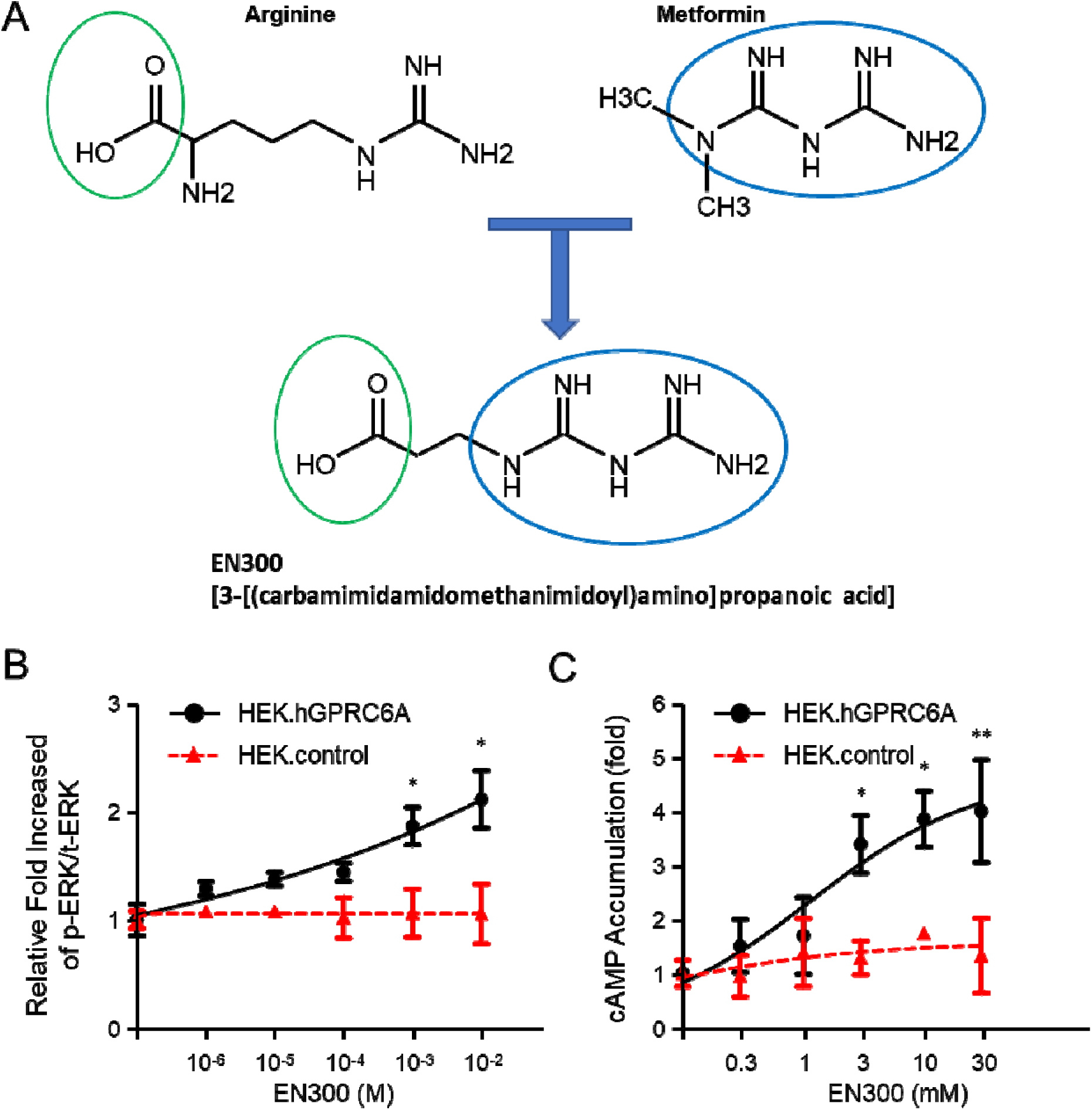
Preliminary structure activity relationships. A. EN300 a with features of both L-Arginine and metformin. B and C. Dose-dependent effects of EN300 on GPRC6A-mediated ERK activation (B) and cAMP accumulation (C). HEK-293 cells were transfected with cDNA plasmids of GPRC6A for 48 hours, after incubated in Dulbecco’s modified Eagle’s medium/F-12 containing 0.1% bovine serum albumin quiescence media for 4 hours, then exposed EN300 at indicated concentrations for 15 minutes for ERK activation, or 40 minutes for cAMP accumulation details as described under “Methods”. *P<0.05, **P<0.001 that indicates a significant difference from control and stimulation groups.

### GPRC6A modifies metformin effects on glucose metabolism *in vivo*

Finally, to assess if GPRC6A impacts metformin’s *in vivo* effects on glucose metabolism, we assessed the effects of metformin to improve glucose control *in vivo* in wild-type and Gprc6a null mice. For these studies weight-matched wild-type and *Gprc6a^-/-^* female mice were fed with normal chow and HFD for 4 weeks, and then treated with vehicle (saline, 10 μl/g) or metformin at dose of 4 mg/g daily i.p. injection for 5 days. Treatment of wild-type mice with HFD resulted in impaired glucose tolerance. Metformin treatment of wild-type mice resulted in lowering of the blood glucose concentrations and AUC (Figure 6A), with the magnitude of the response more striking in HFD feed mice, consistent with metformin’s known effects to lower glucose. In con-trast, the glucose lowering effects of metformin in mice lacking GPRC6A were markedly attenuated in *Gprc6a^-/-^* mice (Figure 6B).

**Figure 6.**
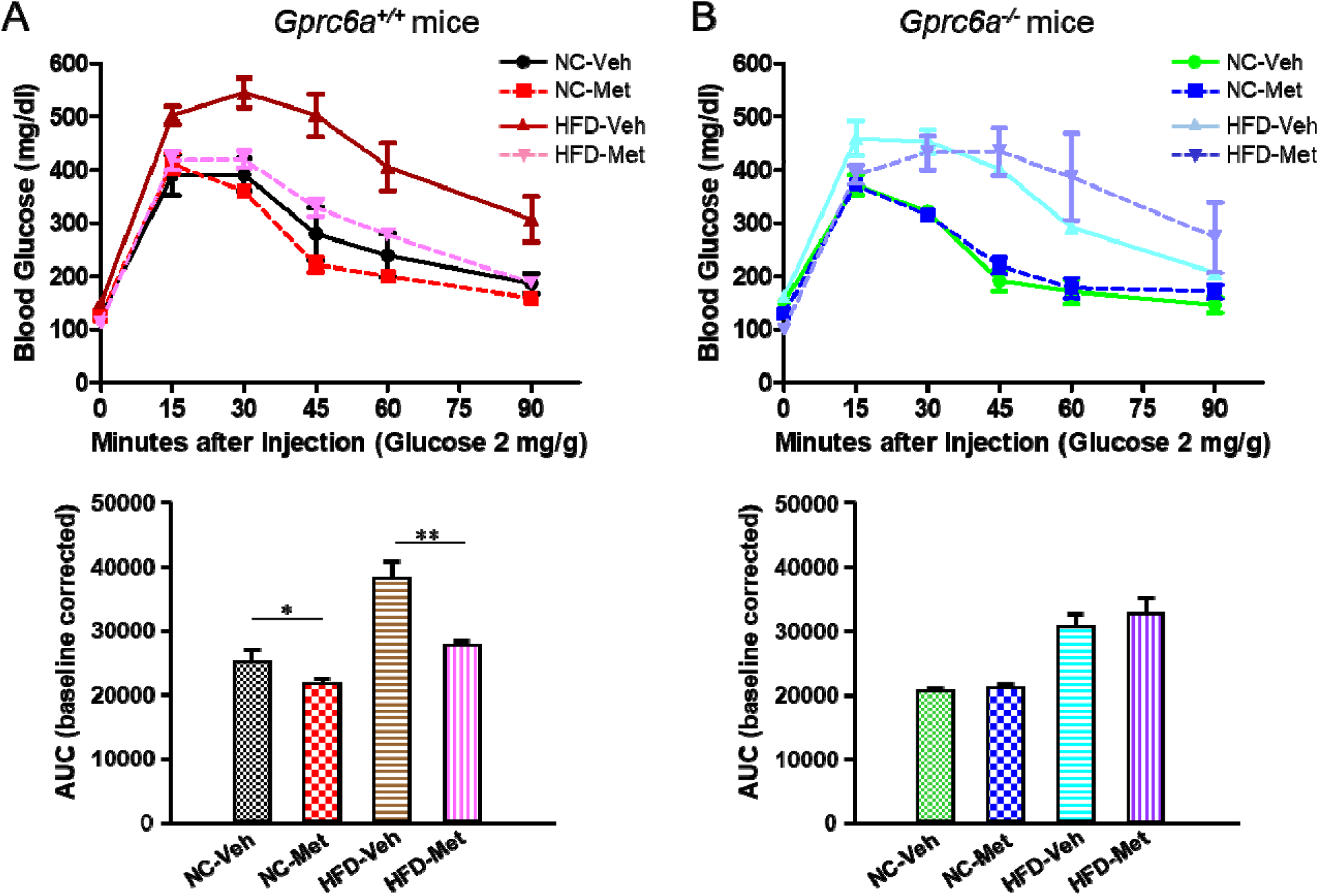
Effects of metformin on glucose tolerance test (GTT) in wild -ype mice (A) and *Gprc6a^-/-^* knockout mice (B) with normal chow (NC) or high fat diet (HFD). 8-week-old female mice were fed normal chow or HFD for 4 weeks, then treated with vehicle (saline; 10 μl/g), or 4 mg/g metformin daily i.p. injection for last 5 days. GTT was performed in mice after 6 hours fasting, mice were treatment with a single i.p. dose of glucose (2 g/kg). The treatment groups are: Normal chow with vehicle (NC-Veh); Normal chow with metformin (NC-Met); High fat diet with vehicle (HFD-Veh); and High fat diet with metformin (HFD-Met). Area under the curve (AUC) for glucose was calculated using the trapezoidal rule is shown on the bottom. Results are means + SE. *P<0.05 and **P<0.001 that indicate a significant difference from vehicle group vs. metformin treatment group in normal chow or HFD.

## Discussion

Using structures derived from Alphafold2 and modeling we characterized the orthosteric binding site for the orthosteric ligand, L-Arginine in GPRC6A. We found that the binding site for L-Arginine was located at the Ser149, Glu170, Ser171, Thr172, and Asp303 in the VFT. Similarly, the other amino acid ligands for GPRC6A, L-Lysine and L-Ornithine, are predicted to bind to the same pocket as L-Arginine in GPRC6A (Supplemental data: Figure S1A and B, Table S1). In contrast, the GPRC6A binding pocket for L-Tryptophan, the orthosteric ligand for the closely related receptor CASR, is too small to accommodate the large aromatic tryptophan side chain. This is consistent with the inability of tryptophan to activate GPRC6A *in vitro* functional assays (37). Thus, the characteristic of the binding pocket defines the amino acid specificity.

Most importantly, we show for the first time that GPRC6A is a potentially new therapeutic target for metformin. We explored this question because of the chemical structural similarities between metformin and L-Arginine and the observation that metformin’s effects to regulate glucose homeostasis overlaps the metabolic phenotype observed with GPRC6A activation. Metformin dose-dependently activates ERK and cAMP signaling in HEK cells transfected with a human GPRC6A and metformin administration improved glucose tolerance in wild-type mice but in GPRC6A knockout mice. We found that metformin could be docked to the orthosteric site in VFT of GPRC6A where L-Arginine binds. This binding site for metformin was confirmed by showing loss of *in vitro* activity in alternatively spliced GPRC6A isoforms lacking this putative pocket. The specific amino acids in GPRC6A mediating the contact with metformin were confirmed by showing that site-directed mutagenesis of essential glutamic acid Glu170 (E170) and aspartic acid Asp303 (D303) residues in the predicted binding pocket resulted in loss of metformin stimulation of GPRC6A *in vitro*.

Finally, we tested EN300, a commercially available compound that has carboxylic acid of L-Arginine attached to the guanidine of metformin. This chimeric molecule also activated GPRC6A. Additional docking studies showed that EN300 bound in the same binding pocket with similar contacts to L-Arg and metformin (SI Figure S2). We have previously shown that gallic acid, which consists of a carboxyl moiety linked to a benzyne ring, is a ligand for GPRC6A that binds to Ser149, Thr172 and Asp303 (73), suggesting that it potentially also occupies a site in the VFT overlap binding sites for L-Arginine and metformin. These findings suggest the structural model of GPRC6A can inform future discovery and optimization of novel small molecules based on the metformin or other chemical scaffolds that bind to the predicted VFT orthosteric site. This could lead to developing metformin analogues with improved potency and specificity.

Our studies are limited by several factors. First, we lack an experimentally validated 3D structure such as cryo-EM of GPRC6A alone or in combination with L-Arginine or metformin. However, previous work confirming the Ocn binding site gives us confidence on using the AlphaFold model for this study. Experimental structural data are needed to confirm our predictions. Second, our assessment of GPRC6A role in mediating metformin’s effects need to be confirmed in long-term preclinical studies. More extensive characterization of the metabolic effects of metformin in GPRC6A null mice should help elucidate the importance GPRC6A dependent and independent metformin targets. Since GPRC6A regulates glucose homeostasis, it is also possible that the observed reduction of metformin effects in Gprc6a^-/-^ mice could result from cross talk with regulatory pathways shared between metformin and GPRC6A. Nonetheless, our findings may help define metformin’s molecular mechanisms of regulate energy metabolism.

Regardless, these studies point to the potential importance of GPRC6A as a therapeutic target for treating metabolic syndrome an T2D and justifies testing pharmacological approaches to activate GPRC6A in preclinical models of obesity and T2D.

## ACKNOWLEDGEMENTS

This work was supported by grants from the National Institutes of Health, NIAMS grant number 1R01AR071930 and NIDDK grant numbers 1R01DK121132 and 1R01DK120567 (LDQ).

The content is solely the responsibility of the authors and does not necessarily represent the official views of the National Institutes of Health.

## AUTHOR CONTRIBUTIONS

Rupesh Agarwal, Micholas Dean Smith and Jeremy C. Smith were responsible for the computational studies, and Min Pi and L. Darryl Quarles were responsible for proposing the underlying premise and the experimental studies regarding GPRC6A. All authors contributed to the writing of the paper.

## Supplemental Figures and Table

**Supplemental Figure 1.**
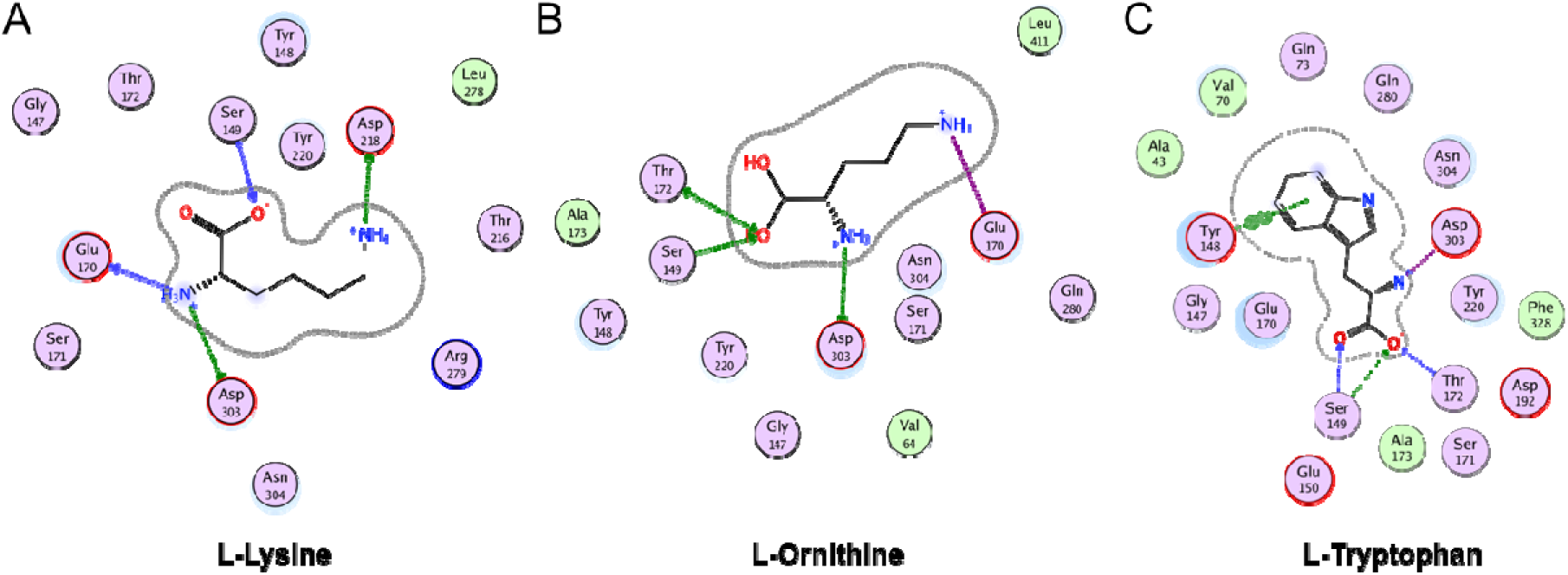
2D ligand-protein interaction maps of L-amino acid ligands for GPRC6A. Docking of L-Lysine (A), L-Ornithine (B) and L-Tryptophan (C), to VFT domain showing residues in binding pocket.

**Supplemental Figure 2.**
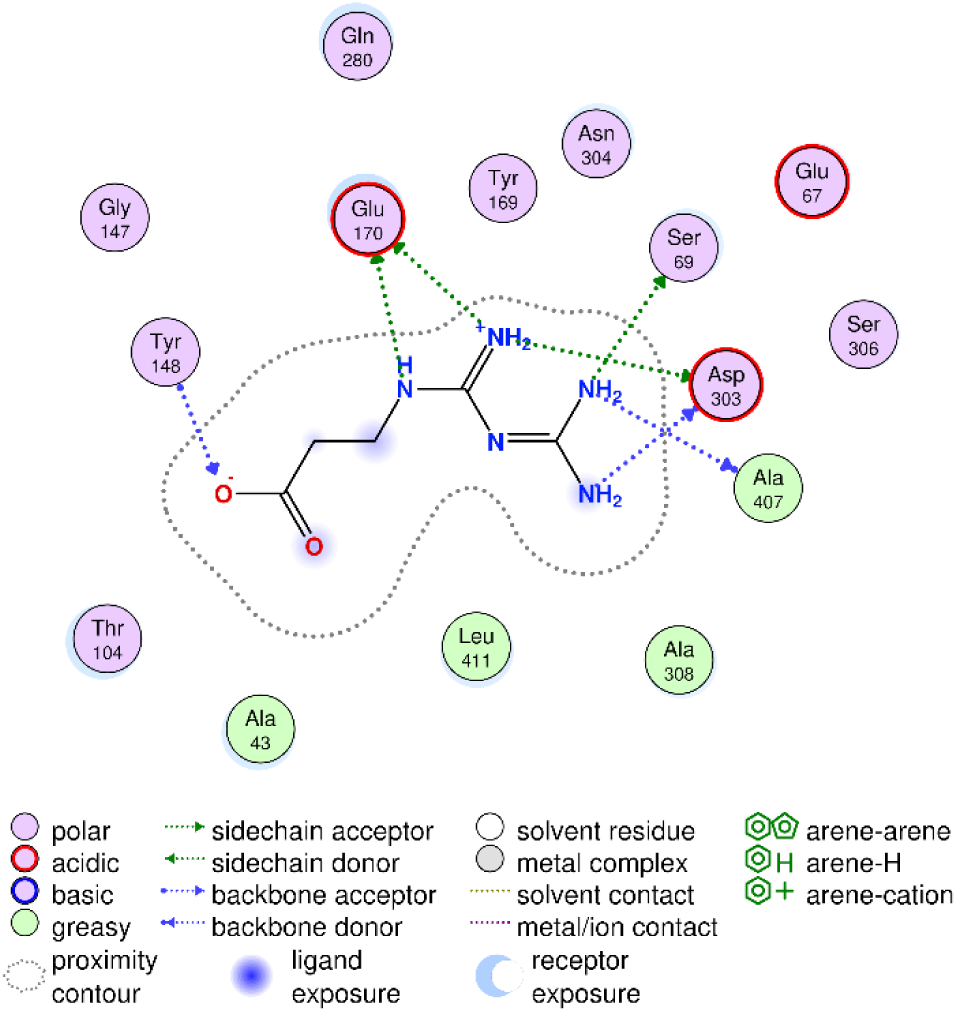
2D ligand-protein interaction map of EN300 for GPRC6A.

**Supplemental Table 1.** Predicted Binding Residues in VFT of GPRC6A.

| GPRC6A<br>position | Ligand |  |  |  |  |  |
| --- | --- | --- | --- | --- | --- | --- |
|  | Arginine | Lysine | Ornithine | Metformin | Tryptophan | EN300 |
| Tyr 148 |  |  |  |  | + | + |
| Ser 149 | + | + | + |  | + |  |
| Tyr 169 |  |  |  | + |  |  |
| Glu 170 | + | + | + | + |  | + |
| Ser 171 | + |  |  |  |  |  |
| Thr 172 | + |  | + |  | + |  |
| Glu 280 |  |  |  | + |  |  |
| Asp 218 |  | + |  |  |  |  |
| Asp 303 | + | + | + | + | + | + |
| Asn 304 |  |  |  | + |  |  |
| Ala 308 |  |  |  | + |  |  |
| Leu 411 |  |  |  | + |  |  |

